# *In vitro* Anti-Cancer Activity of *Oliveria decumbens* Vent. Extract, an Endemic Persian Medicinal Plant, on HT-29 Colorectal Cancer Cell Line

**DOI:** 10.1101/2021.10.01.462701

**Authors:** Amir Khodavirdipour, Fatemeh Haddadi, Hamideh Rouhani nejad, Yasoub Shiri, Veronica Preetha Tilak

**Author notes:** **Corresponding author:** Fatemeh Haddadi, Ph.D. **Address:** Department of Biology, Faculty of Sciences, University of Zabol, 98613-35856, Zabol, Iran **Email:**.

## Abstract

**Introduction:** The top 3 causes of death worldwide include heart disease, injury, and cancer; and cancer records the 2^nd^ place as the leading cause of death in the United States of America after cardiovascular diseases and injuries. Cancer can begin and progress in a very highly twisted and complex pattern and follow the multifactorial route. There is only very few research on medicinal properties *Oliveria decumbens* rare and valuable plant specially on cancer. So, in this study we tried to cover all needs for future *in vivo* research.

**Methods:** MTT assay has been performed to estimate the cytotoxicity of the ethanolic extract of the plant. Its free radical capacity evaluation was done by DPPH assay. Furthermore, real-time PCR, the wound-healing assay along with a DNA damage test to study DNA fragmentation characteristics. The plant’s transcriptomic study was performed by NGS *de Novo* assembly.

**Result:** *Oliveria decumbens* ethanolic extract showed an Ic_50_ of 14.39 μg/ml. The real-time PCR showed that *Oliveria decumbens* ethanolic extract significantly induced apoptosis by upregulating the *bax* gene and slight downregulation of *bcl2* an anti-apoptosis gene. The NGS *de Novo* transcriptome analysis discovered 38 genes responsible for secondary metabolite synthesis so far. The remaining genes and reconstruction of the co-expression network of the transcriptome are underway.

**Conclusion:** The outcome of the Scratch-test and DNA fragmentation confirmed the anti-metastatic and DNA damage properties respectively. Based on these findings; *Oliveria decumbens* ethanolic extract shall be considered as potential anticancer and chemotherapeutic agents which may elucidate in upcoming studies.

## 1 Introduction

### 1.1 Background

Cancer is one of the leading causes of death worldwide according to the WHO (World Health Organization). Out of all types of cancers, colorectal cancer is ranked 3^rd^ after lung and breast cancers and ahead of the prostate, non-melanoma skin cancer, and gastric cancers (ranking produced based on data for both sexes). Unfortunately based on data released by the Iranian ministry of health the growth rate of cancer is almost double the global rate, and gastrointestinal cancers ranked 2^nd^ in the Iranian population considering both sexes. The conventional methods in cancer therapy are surgery, chemotherapy, radiotherapy, and more recently immunotherapy, gene therapy, and near-future CRISPR/Cas9-based therapies. One of the modern therapeutic alternatives in chemotherapy is the use of bacterial extract against cancer cells that are much easier and cheaper to produce in a bio-reactor, one of the latest achievements in the field of bacterial-mediate-apoptosis reported by Khodavirdipour *et al* that showed the effectiveness of *Shigella flexneri* against pancreatic cancer cell line (Khodavirdipour et al., 2020a). Khodavirdipour *et al* in another bacterial-mediated-apoptosis project reported that *Salmonella typhi* has an interesting capacity to induce apoptosis in the same pancreatic cancer cell line (Khodavirdipour et al., 2019b). One of the key characteristics of cancer cells is that they lack proper apoptosis machinery, therefore it can be Achilles’ heel to overcome cancer. *bcl2* gene families including *bax* and *bcl2* are key players in these processes.

Plant-based medicinal by-products are phytochemicals used as therapeutic agents where their usage dates back to early civilization. In the last decades with the advances in biomedicine, molecular genetics and molecular pharmaceutics four categories of plant derivatives chemotherapeutic agents are marketed worldwide including a) the Alkaloids eg. Vinblastine, b) the Epipodophyllotoxins eg. Etoposide, c) the Taxanes eg. Paclitaxel and d) the Camptothecin derivatives eg. Camptothecin. Many other medicinal compounds have the potential to be future safer chemotherapeutic agents. Compounds like carvacrol, thymol, gamma-terpinene, para-cymene, and many others. Many studies conducted and numerous ways are evaluating anti-cancer properties of plant-derived extracts and compounds have proven their pro-apoptotic nature. For instance, Bar-Shalom *et al*., (Bar-Shalom et al., 2019) stated that the extract of *Inula viscosa* inhibits the growth of cancerous cells of colon cancer *in vitro* and also *in vivo*. Nallathambi *et al*. announced that cannabis-derived compounds have an anti-proliferative effect on colon cancer cell lines by cell cycle arrest at S or G0/G1 phases of interphase; in another experiment on HT-29 colorectal cancer cell line (Nallathambi et al., 2018). Khodavirdipour *et al*. stated that the ethanolic extract of *Syzygium cumini* significantly induced apoptosis in the said cell line (Khodavirdipour et al., 2020b). In the other research, the effect of methanolic extract of *Coleus amboinicus* (Mexican mint) by upregulation of *Bax* which is a pro-apoptotic gene, and also activated caspases pathways was shown (Laila et al., 2020). The anti-cancer effects of bitter apricot on pancreatic cancer cells showed that *bax/bcl2* ratio drastically increased and this elucidated that the *bax* gene upregulated and caused induction of apoptosis (Aamazadeh et al., 2020). In another study conducted by Khurshid *et al*., (Khurshid et al., 2020) from Irvine, USA, they reported that *Nigella sativa* extract has an antiproliferative effect on MCF-7 breast cancer cell lines. For the current study, the ethanolic extract of *Oliveria decumbens* Vent. has been subjected to the HT-29 colorectal cancer cell line. *Oliveria decumbens* Vent. is an endemic plant of Iran, and for centuries used as antipyretic, antidiarrheal, abdominal bloating, and generally dyspepsia. This medicinal plant belongs to the Apiaceae family which many members of this family have found out to have anti-cancer effects. In other similar studies; Wu and colleagues stated that the extract of *Angelica sinesis* can inhibit the immunomodulatory function of lung cancer cell lines and down-regulate the expression of *NFκB, STAT3, HIF-1α*, and *VEGF* in tumors (Wu et al., 2019). In another project on members of the Apiaceae family, Goodarzi and researchers evaluated the pro-apoptotic effect of *Cuminum cyminum* on breast cancer cell line (MCF-7) and reported that it can be considered as a future chemotherapeutic drug and further studies has been suggested (Goodarzi et al., 2020).

### 1.2 Aim

Apiaceae family especially *Oliveria decumbens* Vent. are rich in chemical compounds such as carvacrol, thymol, gamma-terpinene, and para cymene. Each of the latter individually has anticancer, anti-proliferative, and pro-apoptotic properties, and interestingly these compounds can be found abundantly in *Oliveria decumbens* Vent. as per our knowledge, this is the first study of this kind. In this study, anti-cancer and pro-apoptotic effects of ethanolic extract of *Oliveria decumbens* Vent. in colorectal cancer studied and meanwhile whole transcriptome and *de novo* assembly of aerial part of *Oliveria decumbens* Vent. was performed and responsible genes for secondary metabolites have been reported and the rest of the transcriptome analysis will be reporting in the next paper.

## 2 Material & Methods

### 2.1 Cell line and culture

ATCC certified Human colorectal cancer cell line HT-29 with serial No.: 300215, has been purchased from the Iranian Biological Resource Centre (IBRC), Tehran, Iran. Cells were cultured in RMPI-1640 with FBS (10%), 100 U/mL of Penicillin in a humidified atmosphere at 37ºC, and 50 μg/mL CO_2_ (5%). Cell viability ascertained by trypan blue dye staining, consequently plated with higher than 90% of viable cells used for the research. Morphological studies of cultured cell lines were performed by an inverted microscope (Khodavirdipour et al., 2019a).

### 2.2 Plant material

Flowers of the plant cultivated at the end of May 2018 from heights of 2000 m of Fars province, collected and identified by Dr. A.R. Khosravi, Department of Biology, Shiraz University-Fars province (herbarium #55075).

### 2.3 Ethanolic extract of *Oliveria decumbens* Vent

Solowey et al. named this method as the gold standard in the evaluation of medicinal plants for their properties, so, it has been chosen as no heat/boiling involve (Solowey, E., Lichtenstein, M., Sallon, S., Paavilainen, H., Solowey, E. and Lorberboum-Galski, H., 2014. Evaluating medicinal plants for anticancer activity. *The Scientific World Journal, 2014*). In this procedure for every 20 g of dried plant Powdered 300 mL of 80% ethanol has been used for six days and filtered by watmann filer-paper, subsequently kept at 37ºC incubator for complete evaporation. For stock solution preparation 0.5 g of extract has been taken into 10 mL of RMPI-1640 culture medium solution. It is necessary to cover the solution with aluminum foil to prevent oxidation.

### 2.4 Essential oil isolation

Powdered flowers were exposed to hydro-distillation for 4h by the Clevenger apparatus. The resulting essential oil was dried by anhydrous sodium sulfate and kept at 4ºC.

### 2.5 Cell viability assessment using MTT assay

Cell viability and proliferative activity of *Oliveria decumbens*Vent. treated HT-29 cell line was evaluated using 3-(4,5-dimethylthiazol-2-yl)-2, 5-diphenyl-tetrazolium bromide (MTT) which examines the percentage of viable cells (Khodavirdipour et al., 2021). The HT-29 cell line with 75% convergence was separated from the plate by 0.05% EDTA-trypsin. Cells were transferred into 96 well plates with a concentration of 10^4^ cells. Three wells were kept as untreated or control. The assay was carried out for 24 hours after treatment with different concentrations of *Oliveria decumbens* ethanolic extract such as 2, 4, 6, 8, 10, 12, 14, 16, and 18 μL/mL. To prepare 3-(4, 5-dimethylthiazol-2-yl)-2, 5-diphenyl-tetrazolium bromide (Sigma-Aldrich) powder, 2 mg of said reagent dissolved in 1 mL PBS. The medium was substituted with 180 μL of freshly prepared media along with MTT reagent, untreated wells considered as control. Plate incubated at 37ºC with 5% CO_2_ and sufficient humidity for 4h; then after discarding the solution of the well, 150 μL of 2-propanol was added to each well. To prevent light from reacting with samples it is covered with aluminum foil and shake at 100rpm for 15min. Optical density was read by an ELISA reader at 570nm. The test was performed three times for greater accuracy for each concentration of the extract and the control, finally, IC50 of each concentration was recorded (He et al., 2016; Neubig et al., 2003).

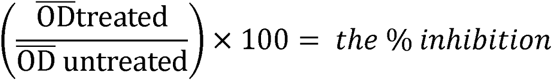

Since the commercialization of MTT assay (colorimetric technique)-this method has performed widely to determine IC50 in a much faster and easier path.

### 2.6 DPPH assay

DPPH assay is the common abbreviation for the 2, 2-diphenyl-1-picrylhydrazyl organic chemical compound test. 0.5 ml of different concentrations of *Oliveria decumbens* ethanolic extract added to 0.5 ml of DPPH solution. This is 0.1 mM in ethanol. The mixture was incubated for half an hour at room temperature. OD was read at 517 nm. The scavenging activity of each concentration compared to the blank. Ascorbic acid is also employed as a control. The following formula used to calculate the anti-oxidant activity of the extract:

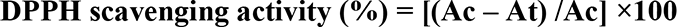

Where Ac is the absorbance of the control and at is the absorbance of the test sample (Shisode and Kareppa, 2011).

### 2.7 RNA extraction and cDNA synthesis

Cultivated HT-29 cells were treated with 14.39 μg/mL of *Oliveria decumbens* for 24h followed by. RNA extraction using the RNX-Plus kit (CinnaGen, IRAN) according to the manufacturer‘s instruction. The quality and quantity of extracted RNA were evaluated by Nanodrop 1000 spectrophotometer (Thermo scientific-USA) at 260/280 ratio.

cDNA synthesis was performed using Takara-JAPAN based on manufacturer protocol and synthesized cDNA was stored at -20°C until use.

### 2.8 Real-time PCR technique

Real-time PCR was used to evaluate the changes in the expression of *Bax, Bcl2*, and β*-actin* and their fold change as apoptosis-associated genes. HT-29 cells were cultured in 96 well plates in presence of 14.39 μg/mL of ethanolic extract of *Oliveria decumbens* vent for 24h. control cells were treated with DEPC.

RT-PCR was performed using the SYBR green master mix (RealQ plus 2X, Qiagen-Germany). Amplification was done by Exicycler96 (Bioneer-S. Korea) in a total volume of 20 μL. Each well contained 1μL of each reverse and forward primers (Macrogen-S.Korea), 2 μL cDNA, and 10 μL of master mix. Primers used in this experiment and the corresponding annealing temperature of each are shown in table 1.

**Table 1:**
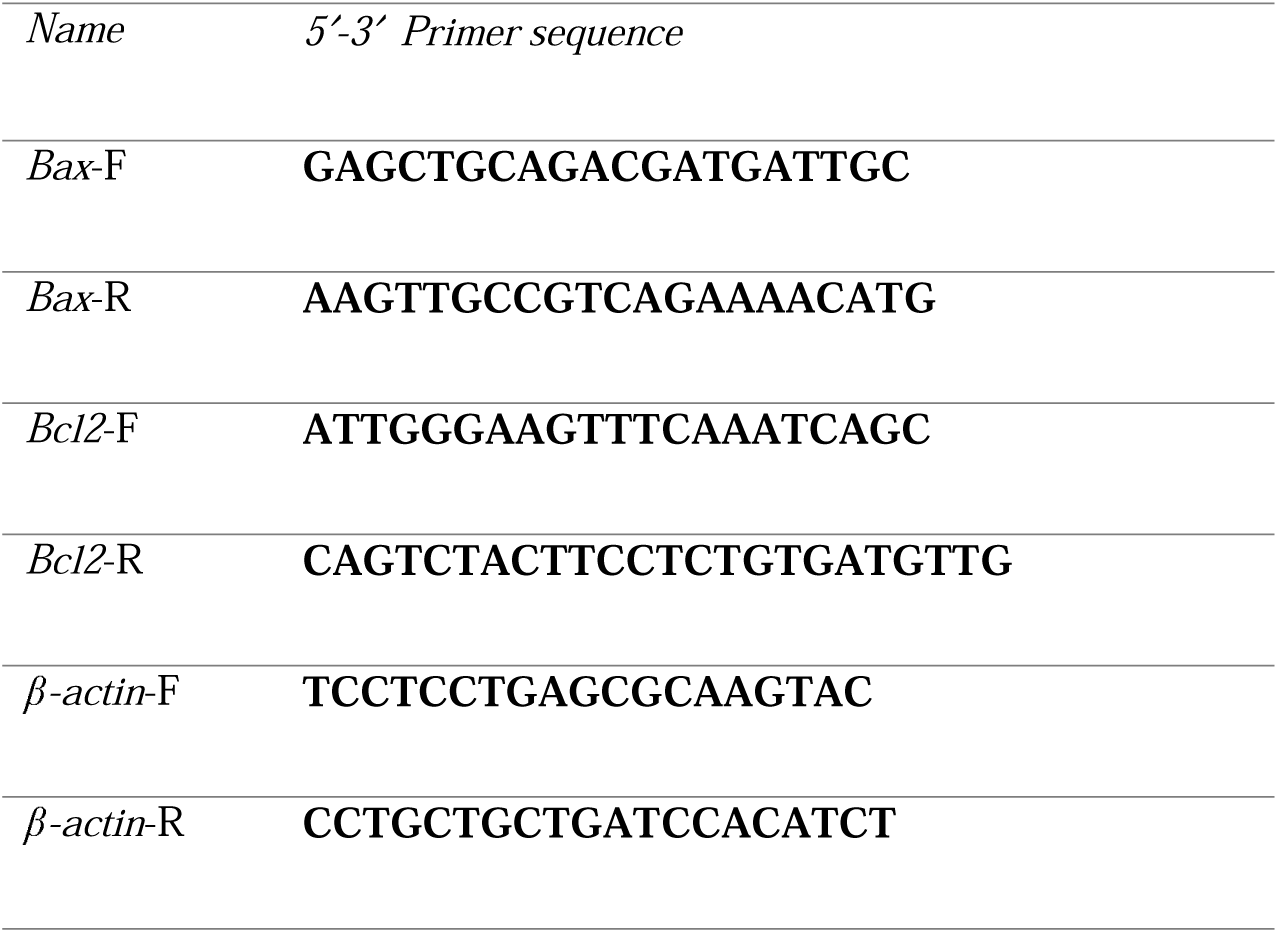
Real-time PCR primers sequence.

### 2.9 RNA-seq

*Oliveria decumbens* Vent. high quality extracted RNA has been sent to Macrogen Co., South Korea in the dry-ice package. Received transcriptome analyzed and annotated for finding the key genes involved in the production of medicinal by-products and secondary metabolites analysis was performed using the Velvet algorithm and CLC genomics workbench.

### 2.10 Scratch-test Assay

The scratch-test (wound healing assay) was performed to evaluate the HT-29 cell-migration ability. Following the procedure cells were placed in 6 well culture plates and allowed to grow and make a merging monolayer; the scratch was created by using a sterile tip of yellow pipette; to remove free-floating cells or any debris cells washed thoroughly. At next, the cells exposed to the ethanolic extract of *Oliveria decumbens* Vent. and cells kept in an incubator at 37 °C for 48 h. the progression of wound healing was assessed at 0, 24, 48, and 72 hrs intervals for scratch line and images recorded at the moment using the light microscope.

### 2.11 DNA Damage Analysis before and after of ethanolic extract of *Oliveria decumbens* Vent

Lastly, a DNA extraction kit (InstaGene Matrix-Bio-Rad – USA) is utilize zed for this purpose. Colorectal cancer cells seeded into 6 wells plate for 24 hours and later exposed to concentration (Ic50) of *Oliveria decumbens* Vent. DNA extracted after 48 hours following the kit’s protocol. Gel electrophoresis was utilized to assess the DNA break.

### 2.12 Statistical analysis

All the obtained data were analyzed by using SPSS v. 25 by considering the P-value < 0.05.

## 3 Results

### 3.1 MTT assay

Based on MTT assay performed on the ethanolic extract of *Oliveria decumbens* vent. We found out that 14.39 μg/mL can expose their toxicity on the human colorectal cancer cell line after 24 hrs. (The test performed in a time and dose-dependent manner of 24, 48, and 72 hrs) after using the Pearson Correlation *Coefficient formula* and obtained data showed *Ic50* at 14.39μg/mL. Calculation of the inhibitory concentration (50% maximal) is the key value for a better and deeper understanding of biological and to be exact pharmacological characteristics of a potential chemotherapy agent.

### 3.2 Cell line treatment with the *Oliveria decumbens* Vent. extract

After the treatment of the HT-29 colorectal cell line with 14.39 μg/mL of *Oliveria decumbens* vent. ethanolic extract increase in apoptosis rate has been observed in cancer cell lines after 24h. Figure 1 is showing the cell line before and after treatment by the ethanolic extract of *Oliveria decumbens* vent. Mean of three times repeat of the test and calculated by the previously stated formula.

**Figure 1:**
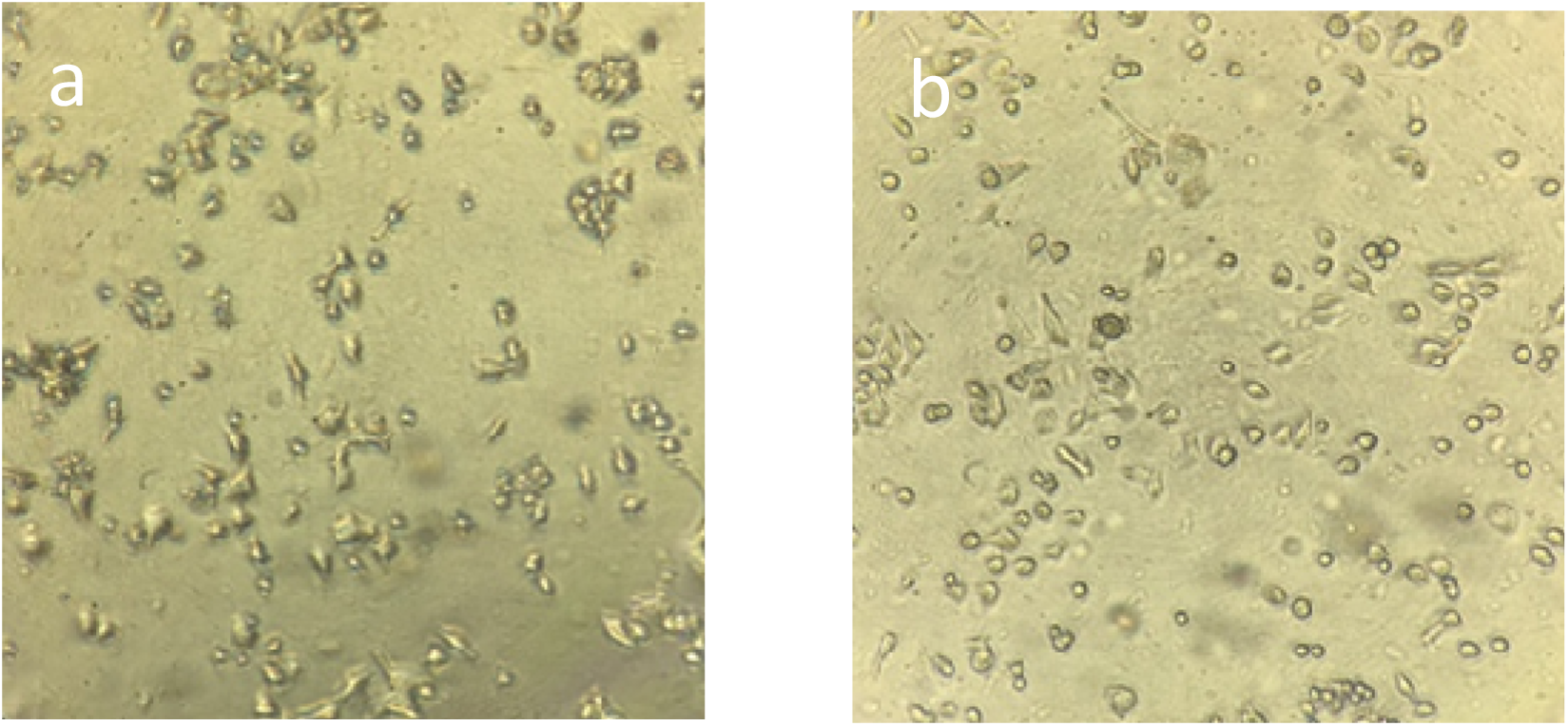
Microscopic view of colorectal cancer cell line. **a)** Control and untreated colorectal cancer cell.

### 3.3 DPPH

As shown in Figure 2, by an increase in the Ethanolic extract of *Oliveria decumbens Vent*. The potency of inhibition of DPPH free radicals also increases and in higher concentration the difference between inhibition potency reduces. (P>0.05).

**Figure 2:**
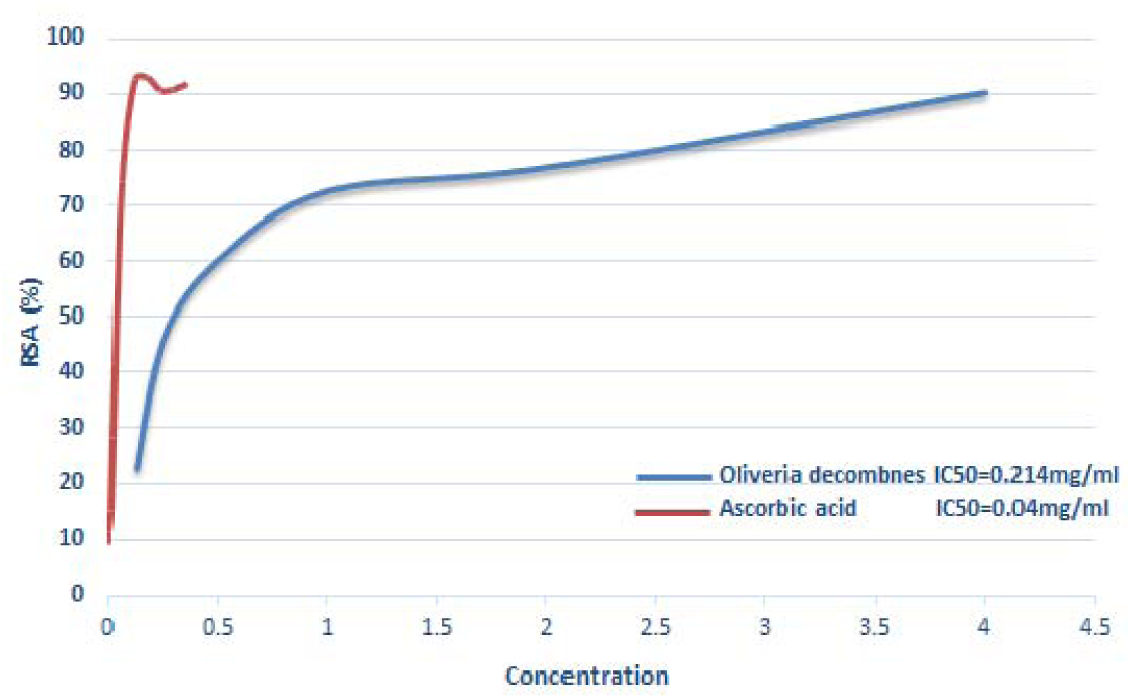
Mean comparison of antioxidant activity of ethanolic extract of *Oliveria decumbens* Vent. vs Ascorbic acid.

Based on obtained data Ic50 from the ethanolic extract of *Oliveria decumbens Vent*. determined as 0.214 mg/mL and for ascorbic acid 0.04 mg/mL. The results suggesting that the ascorbic acid value is much lesser than the value of the ethanolic extract of *Oliveria decumbens Vent*. so ascorbic acid is a much potent antioxidant.

### 3.4 Real-time PCR

Real-time used to analyze the expression level of *Bax* and *Bcl2* genes in the HT-29 cell line before and after treatment with ethanolic extract of *Oliveria decumbens* vent. the quality of total extracted RNA determined by Nanodrop spectrophotometer (Thermo Scientific, USA) RT reaction (Reverse transcriptase and synthesis of cDNA done by cDNA kit (Takara-Japan) based on kit’s protocol. The cDNA product was amplified using real-time PCR (Bioneer-S. Korea) with the mentioned designed primers. The obtained results showed in table 2.

**Table 2:**
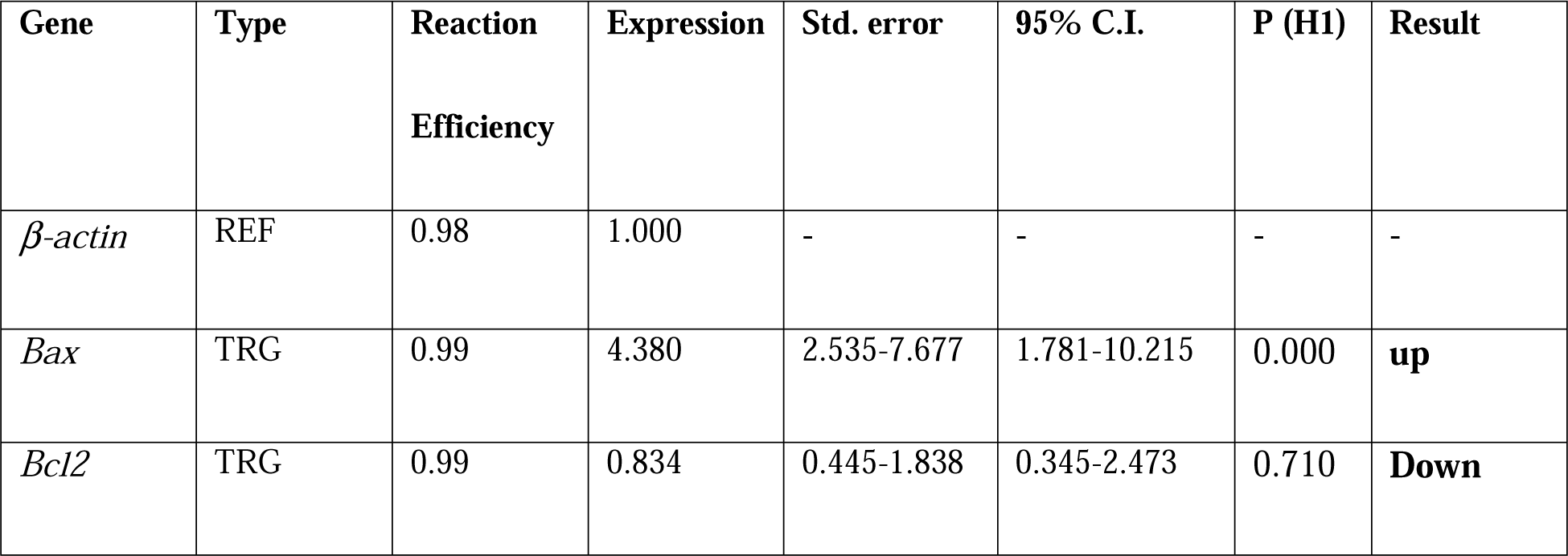
Standardized result of Real-time PCR, *Bax* and *Bcl2* genes expression analysis. Legend: *P(H1) - Probability of alternate hypothesis that difference between sample and control groups is due only to chance. TRG: Target, REF: Reference*.

Based on standardized expression data of the pro-apoptotic gene of *Bax* on the treated cell of HT-29 with 14.39 μg/ml of Ethanolic extract of *Oliveria decumbens* Vent. Shows 4.38 fold increased while the expression of the *Bcl2* gene which is an anti-apoptotic gene down-regulated to 0.8 from 0.99 in comparison to the reference gene (Table 2).

### 3.5 RNA-seq

In table 3 the expression amount of 5 key genes in terpenoid synthesis in Oliveria decumbens Vent. has been mentioned. So, for better understanding, the expression pattern of another 33 genes with different functions is also mentioned in the above table. As it can be observed, genes involved in terpenoid synthesis had more than 3000 expression copies. Cytochrome P450 gene has the highest expression among the 5 genes that are responsible for secondary metabolite synthesis in the plant, and more importantly, that has far different functions and a variety of activities besides the above-mentioned role)McDonnell and Dang, 2013) By considering the said fact, its high expression can purely be justified. Gamma-terpinene synthase enzyme is located at the bullseye point of the terpenoid synthesis pathway and the high expression copy of that guarantees the terpenoids synthesis)Shimada et al., 2004).

**Table 3:**
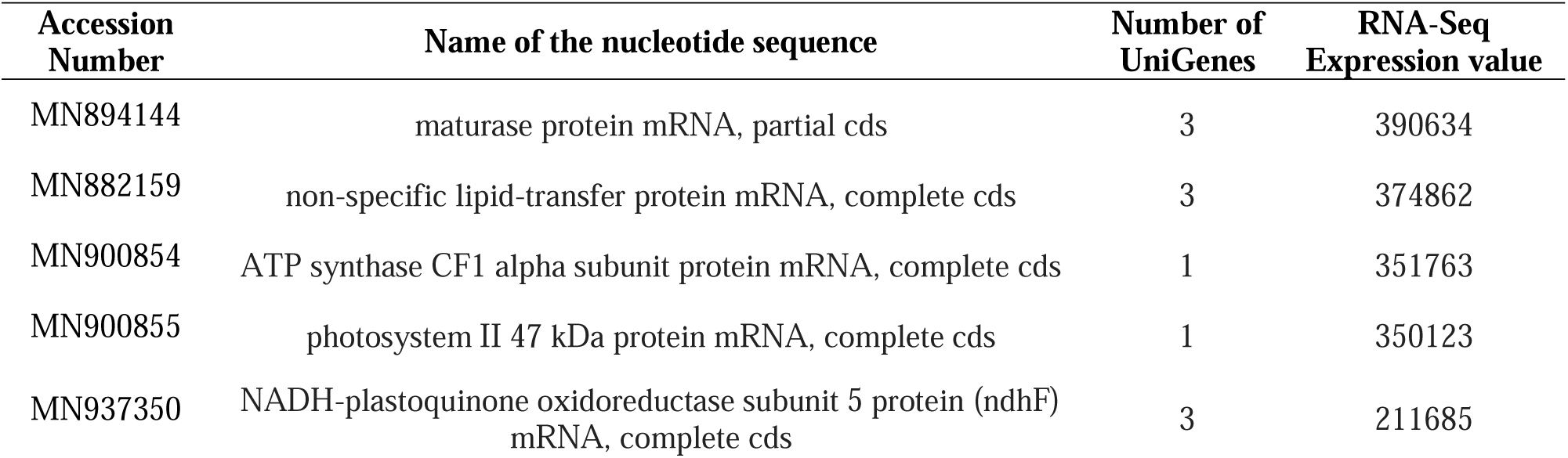
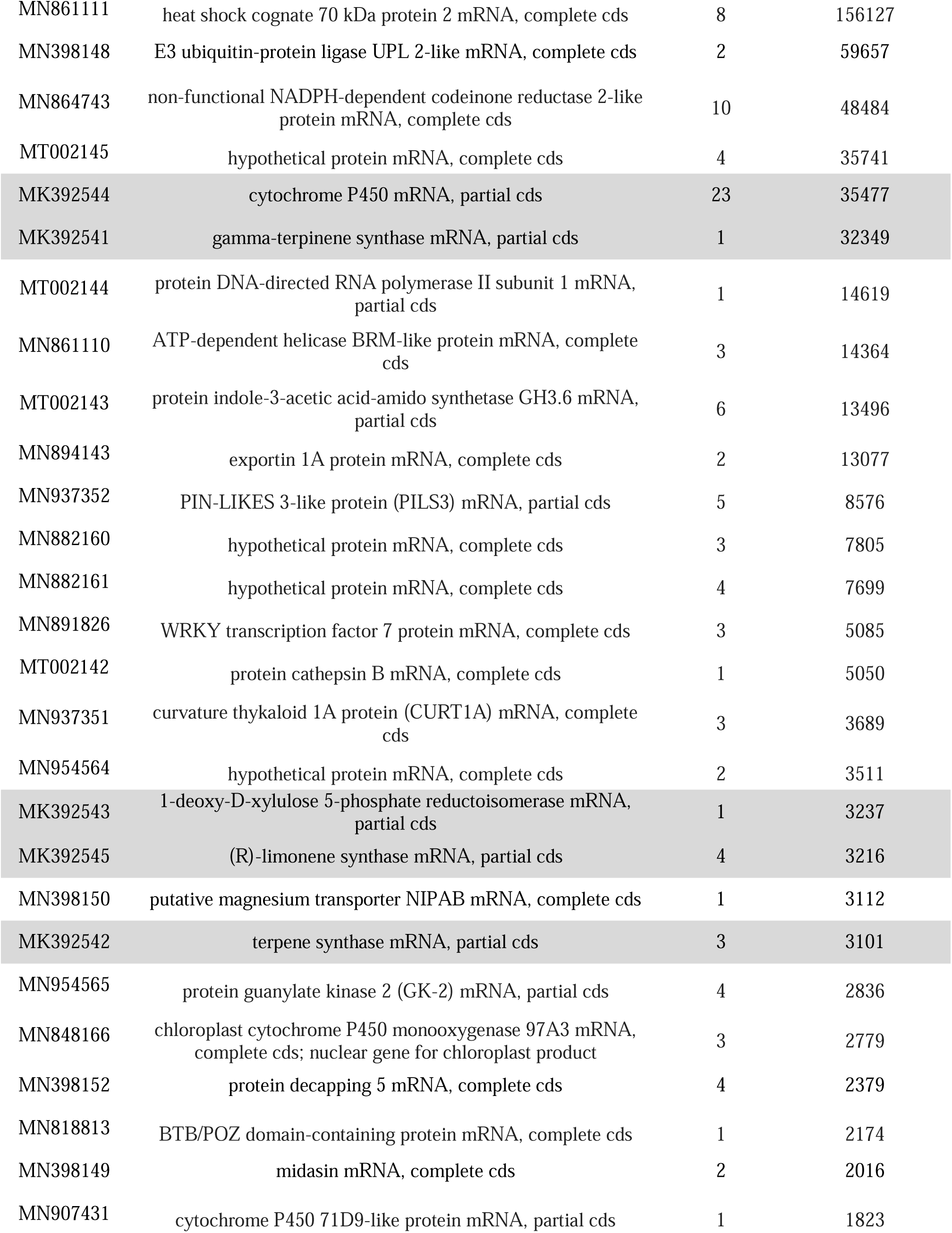
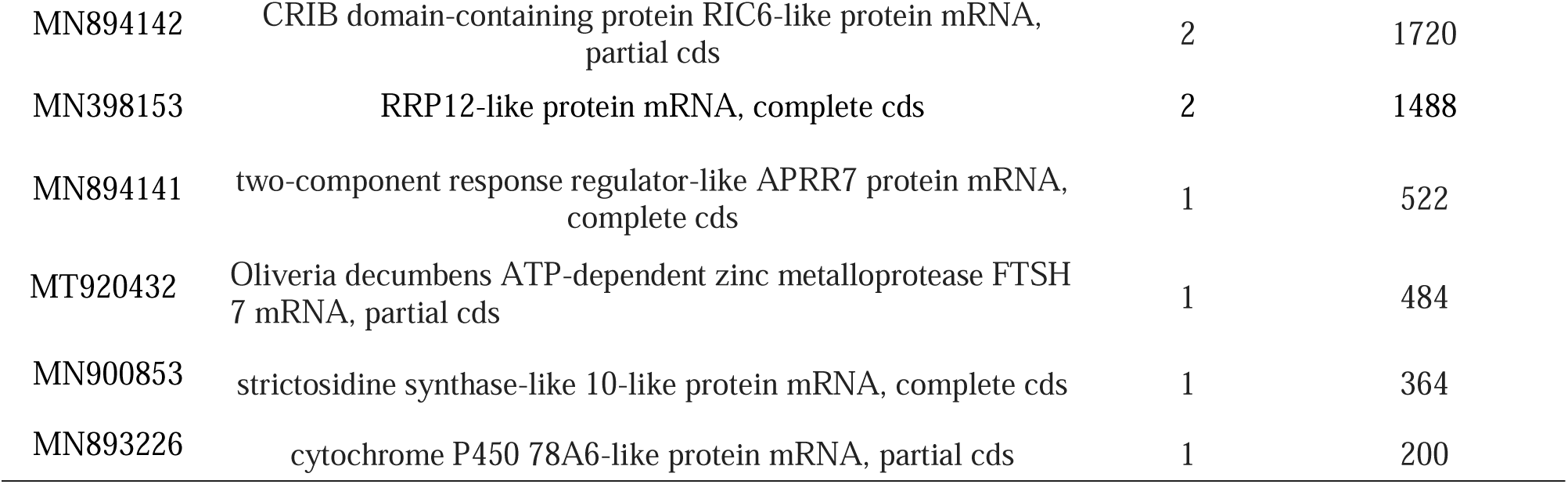
*Oliveria decumbens* Vent. Genes and their corresponding Ac No. at NCBI database. Expression profile of some of known genes involved in terpenoid synthesis in plants marked in grey colore, and the rest showing the expression pattern of another 33 genes with different functions.

The preliminary result of the transcriptome assembly and gene annotation so far identified the first 38 genes responsible for medicinal secondary metabolites which have been deposited to the NCBI database by following Ac. No. (Table 3).

### 3.6 Scratch-test (wound-healing assay)

This test was conducted to assess the effect of the ethanolic extract of *Oliveria decumbens* Vent. on cell motility and migration. The result of the present test showing that the higher conc. of the extract results in lower mobility of the cells and migration and consequently can be nominated as a potential anti-metastatic agent (Figure 3).

**Figure 3:**
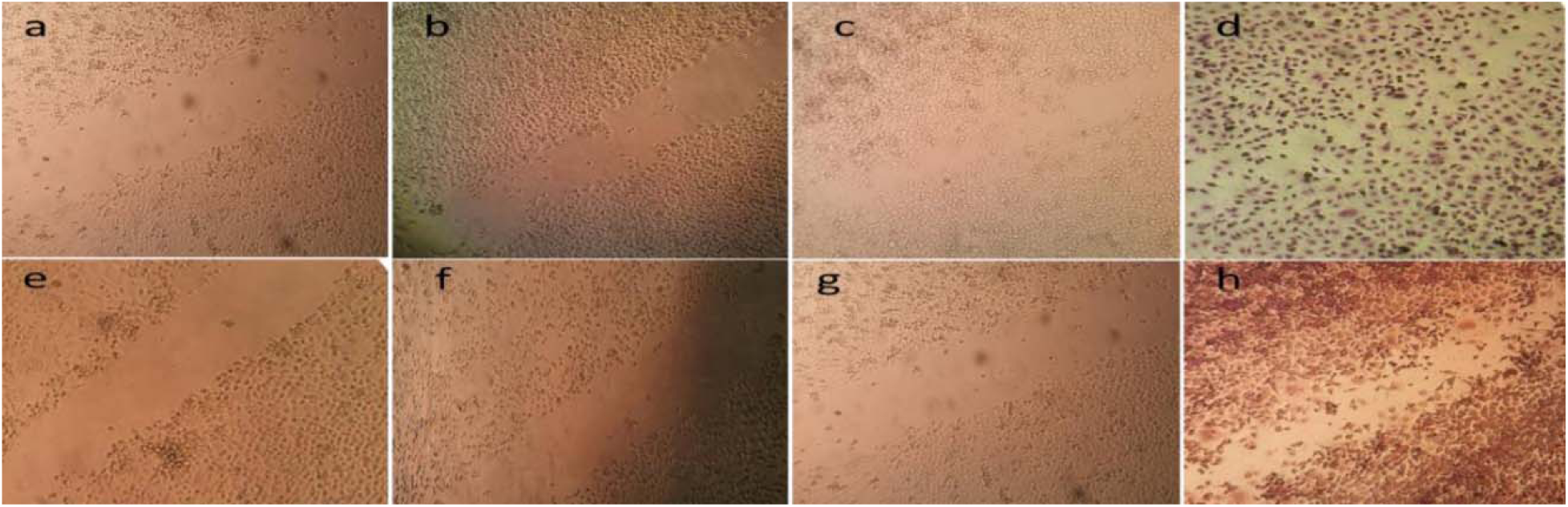
The first row (a-d) showing the time-dependent control from 0 to 72 hours and 2^nd^ row (e-h) showing the cell line treated with ethanolic extract of Oliveria decumbens Vent on the same time frame.

### 3.7 DNA Breaks Assay

The evaluation of the DNA breaks and damage shows that significant DNA damage in HT-29 cells exposed to the ethanolic extract of *Oliveria decumbens* Vent. in comparison to the control group. The smear was observed for the test group while the sharp band formed for the control group (Figure 4).

**Figure 4:**
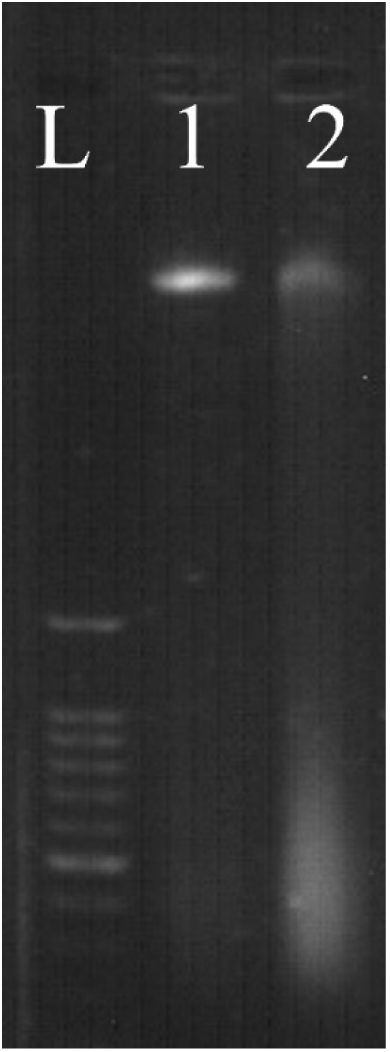
DNA fragmentation and damage after ethanolic extract of *Oliveria decumbens* Vent treatment (1) which DNA smear can be seen clearly; in comparison to untreated control (2). The ladder is marked by (L).

## 4 Discussion

As said before, this is one of the pioneering studies to evaluate the anti-cancer properties of this rare plant in its ethanolic form and more importantly to do NGS *de novo* transcriptome of the plant. The current study is one of its kind, but Lai and colleagues in 2012 in Taiwan reported that Carvacrol and Thymol as a secondary metabolite of many medicinal plants have very low cytotoxicity compared to other medicinal molecules found in plants under their study. Due to the high amount of carvacrol and thymol, P-cymene and Gama-terpinene were reported by Hajimehdipoor et al in *Oliveria decumbens* Vent. we can assume that its low toxicity is because of the presence of the above secondary metabolite (Haji Mahdipour et al., 2010). Regarding the *Oliveria decumbens* Vent. DPPH capacity, in comparison to other Apiaceae family species like *Foeniculum vulgare* Mill. (Fennel) and Trachyspermum *Ammi* L. (carom) which was reported by Goswami and Chatterjee (Goswami and Chatterjee, 2014) *Oliveria decumbens*Vent. has lesser antioxidant activity yet good enough as the daily based antioxidant source. In terms of anti-proliferative and anti-cancer properties, current results showed that the extract directly induced apoptosis by up-regulating the pro-apoptotic gene of *Bax* which is responsible for the induction of cell death. Our findings are in harmony with Wang and the team by experimenting on ginkgo Biloba extract on Liver fibrosis isolated cell by western blotting and immunohistochemistry found that *Bax*:*Bcl2* ratio increased by few fold and the pro-apoptotic gene has been up-regulated (Wang et al., 2015). Deng and co-researcher used pomegranate pill extract on prostate cancer’s metastatic cell line by western blot analysis proven that expression of *Bax* gene significantly increased in comparison to *Bcl2* gene which is down-regulated in their study and also it has been observed that activation of caspase 3 too has a synergic effect on induction of apoptosis beside up-regulation of *Bax* gene (Deng et al., 2017). In a very latest study done by Czerwonka and colleagues, hydro-extract of spirulina algae on A549 cell line of pulmonary cancer demonstrated that spirulina extract by cell cycle arrest on G1 phase induced apoptosis on the said cell line, furthermore this extract inhibits the phosphorylation of Akt, Rb and also decreased expression of cyclin D1 and CDK4 but increased the expression ration of *Bax* to *Bcl2* genes (Czerwonka et al., 2018). In the same year of 2018, Lenzi and team showed that ethanolic extract of *Meripilus giganteus* on the leukemic cell line of HL-60 and Jurkat, by flow cytometry analysis found that this extract elevates the ratio of *Bax* gene in comparison to the anti-apoptotic gene of *Bcl2* and suppress the intercellular ROS and upregulate the expression of Fas. Indeed, ROS level suppression is associated with the activation of the apoptosis mechanism (Lenzi et al., 2018). Besides, *Meripilus giganteus* extract causes cell cycle arrest in the HL-60 cell line. Esmaeili-Mahani and team by the study on *Stureja Khuzestanica* extract on MCF-7 breast cancer cell line in comparison with Vincristine, a chemotherapeutic agent found that after 24 hours and 150 and 200 microgram/ ml of the extract by activation of caspase 3 and increase in the ratio of *Bax*: *Bcl2* induces apoptosis on MCF-7 Breast cancer cell line (Esmaeili-Mahani et al., 2018). Oncologists suggest that now it is the time to emphasize plant-based chemotherapeutic agents and as current invasive procedures such as surgery or conventional treatment like radiotherapy and chemotherapy had limited effect and in some cases gain resistance in cancer cells. Yin and colleagues reported that Carvacrol one of the main medicinal compounds in the Apiaceae family including thymus Vulgaris and *Oliveria decumbens* by activating the mitochondrion pathway and mitogen-activated protein kinase can induce apoptosis in HepG2 hepatocellular carcinoma (Yin et al., 2012). In another study by Fan and co on colon cancer cell lines of HCT-116 and LoVo, carvacrol had an antiproliferative effect and by activating the mitochondrion pathway and also utilizing MAPK and PI3k/Akt pathways induces apoptosis in said cell lines (Fan et al., 2015). Khan and colleagues demonstrated the effects of carvacrol on pulmonary cancer *in vitro* and *in vivo* as well and found that in both situations by carvacrol induced apoptosis by activating mitochondrial pathway (Khan et al., 2018). On the other side, Kang and colleagues evaluated the effects of thymol on gastric carcinoma cells which confirmed induced apoptosis by cell cycle arrest on the G2/M phase (Kang et al., 2016). Anna Marchese and the team in a comprehensive review confirmed that p-cymene has an anti-cancerous effect so besides strong antibacterial effects (Marchese et al., 2017). Zi-li wu and team reported that gamma-terpinene has a strong role in human lung cancer (Wu et al., 2016).

NGS (new generation sequencing) is one the latest technology in molecular genetics that empower us to expand our knowledge about the desired organism in molecular level and it will enable us to find out about any mutation or even help us to identify new genes or maybe identify and report new genus or species. Besides that sometimes we can do WGS (whole genome sequencing) or whole transcriptome analysis for an organism with Refseq or even *de Novo*. In the case of *Oliveria decumbens* as there was not any previous molecular data or Refseq so we opt for Illumina *de Novo* transcriptome sequencing to find out genes responsible for the synthesis of secondary metabolites which had a key role in combating cancer in our *in vitro* study. For our purpose RNA isolation has been performed in our research center and with special packing of dry-ice send to Macrogen-Seoul (S. Korea) for *de Novo* transcriptome sequencing. The received data has been assembled, annotated, and analyzed by the Velvet algorithm, CLC workbench v.20, and with the help of NCBI for alignment and BLAST with *Dacus Carota* as a family member from *Apiaceae* where its whole genome Refseq is available online.

RNA-seq specific *de novo* sequencing for genes involved in secondary metabolite has been received in raw form and analyzed by bioinformatics tools and aligned and BLAST for corresponding genes, the result which has to be 38 genes so far, has been deposited to NCBI database by following Ac. No. (Table 3). The analysis of the entire *de Novo* transcriptome is going on, which will be produced in another paper once completed.

For the first time, Khodavirdipour *et al* in earlier 2020 reported on the anti-metastatic and DNA fragmentation capacity of *Syzygium cumini* and suggested that other medicinal plants may have similar properties which are in total accordance with our findings (Khodavirdipour et al., 2020b).

## 5 Conclusion

By concluding all above can assume that effect of ethanolic extract of *Oliveria decumbens* Vent. by increase induced apoptosis on colorectal cancer of HT-29 by 4.38 fold is in all above reported data and studies as *Oliveria decumbens* has all above medicinal compounds in high percentage and as it has very low toxicity, by enlarge, it will be a great candidate and potential chemotherapeutic agent and anti-cancer drug after *in vivo* and clinical trials.

## Abbreviation

CRISPR/Cas9: clustered regularly interspaced short palindromic repeats/CRISPR-associated complex
DPPH: 2,2-diphenyl-1-picrylhydrazyl
IBRC: Iranian Biological Resource Centre
MTT: 3-(4,5-dimethylthiazol-2-yl)-2,5-diphenyltetrazolium bromide
WHO: World Health Organization

## Declaration

### Ethics approval and consent to participate

Not applicable

### Consent to publish

Not applicable

### Availability of data

All data generated or analyzed during this study are included in this published article. The dataset of 38 genes generated and analyzed during the current study are available in the NCBI repository, are accessible by the mentioned Ac. Numbers in the respective table.

## Authors Contributions

AK – Conceptualization, literature research, performing research, data collection, data analysis, writing manuscript. FH –research supervision, study design, editing manuscript. HR - Research supervision, providing laboratory facilities. YS - Research supervision, data analysis. VPT – Research supervision, editing manuscript. All authors read and approved the final version of the manuscript.

## Acknowledgments

We would like to thank the lab member and supporting staff across both centers in Tehran and Zabol.

---

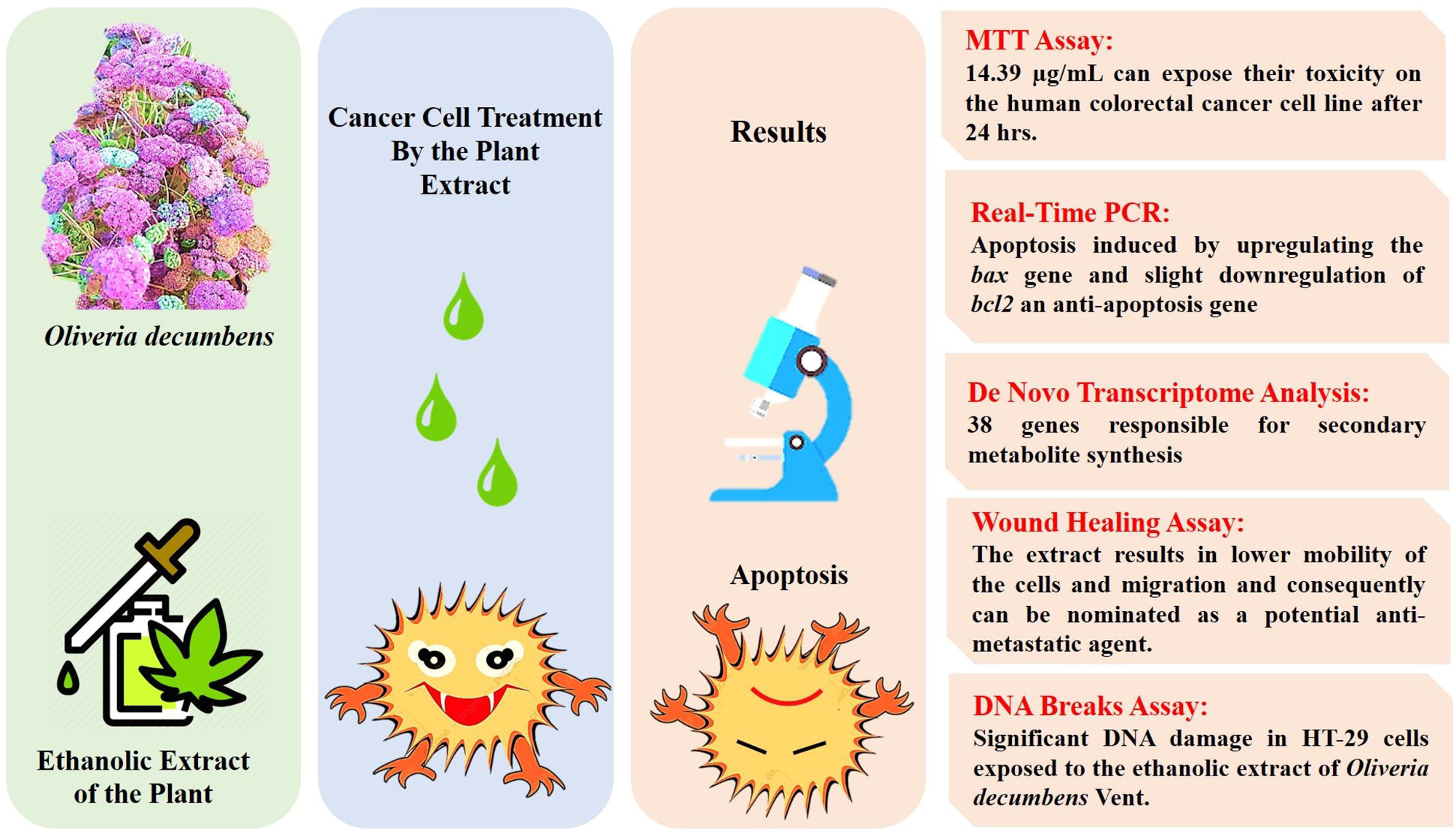

